# Enhancing HBV-specific T cell responses through a combination of epigenetic modulation and immune checkpoint inhibition

**DOI:** 10.1101/2024.09.06.611632

**Authors:** Melanie Urbanek-Quaing, Yin-Han Chou, Manoj Kumar Gupta, Katja Steppich, Birgit Bremer, Hagen Schmaus, Katja Deterding, Benjamin Maasoumy, Heiner Wedemeyer, Cheng-Jian Xu, Anke R M Kraft, Markus Cornberg

**Affiliations:** Department of Gastroenterology, Hepatology, Infectious Diseases and Endocrinology, Hannover Medical School, Carl-Neuberg-Strasse 1, 30625 Hannover, Germany; Centre for Individualised Infection Medicine (CiiM), a joint venture between Helmholtz-Centre for Infection Research and Hannover Medical School, Feodor-Lynen-Strasse 11, 30625 Hannover, Germany; German Center for Infection Research (DZIF), partner site Hannover-Braunschweig, Germany; TWINCORE, Centre of Experimental and Clinical Infection Research, a joint venture between Helmholtz-Centre for Infection Research and Hannover Medical School, Feodor-Lynen-Strasse 7, 30625 Hannover, Germany; Cluster of Excellence RESIST (EXC2155), Hannover Medical School, Carl-Neuberg-Strasse 1, 30625 Hannover, Germany

**Keywords:** hepatitis B virus infection, CD4 T cells, CD8 T cells, T cell exhaustion, checkpoint inhibition, anti-PD-L1, decitabine, DNA methyltransferase inhibitor, epigenetics, HBcrAg

## Abstract

**Objective:** Chronic hepatitis B virus (HBV) infection results in the exhaustion of HBV-specific T cells and the development of epigenetic imprints that impair immune responses and limit the effectiveness of immune checkpoint inhibitor (ICI) monotherapy, such as αPD-L1. This study aimed to determine whether the DNA methyltransferase inhibitor decitabine (DAC) can reverse these epigenetic imprints and enhance the efficacy of ICI in restoring HBV-specific T cell responses.

**Methods:** We investigated HBV-specific CD4^+^ and CD8^+^ T cell responses by 10-day *in vitro* stimulation of peripheral blood mononuclear cells (PBMCs) from patients with chronic HBV infection. PBMCs were stimulated with HBV core-specific overlapping peptide pools and HLA-A*02-restricted peptides, including core_18_ and pol_455_. The immunomodulatory effect of the combination of DAC/αPD-L1 was assessed via flow cytometry. Responder stratification was investigated by comparison of clinical characteristics, *ex vivo* DNA methylation analysis of PBMCs, and determination of IFNγ plasma levels.

**Results:** Treatment with DAC and αPD-L1 enhanced HBV-specific CD4^+^ T cell responses in a significant proportion of 53 patients, albeit with variability. The effect was independent of the HBcrAg level. *Ex vivo* DNA methylation revealed hypermethylation of key genes like *IFNG* among DAC-responders versus non-responders, supported by altered *ex vivo* IFNγ plasma levels. Further analysis of HBV-specific CD8^+^ T cell responses in 22 HLA-A*02-positive patients indicated distinct response patterns between HBV-core_18_- and HBV-pol_455_-specific T cells, with pol_455_-specific CD8^+^ T cells showing increased susceptibility to DAC/αPD-L1, surpassing αPD-L1 monotherapy response.

**Conclusions:** The combination of DAC and αPD-L1 shows promising effects in improving HBV-specific T cell responses *in vitro*. Our study highlights the potential of remodeling exhaustion-associated epigenetic signatures to enhance HBV-specific T cell restoration, suggesting a novel immunotherapeutic avenue for chronic HBV infections.

**Graphic Abstract:** 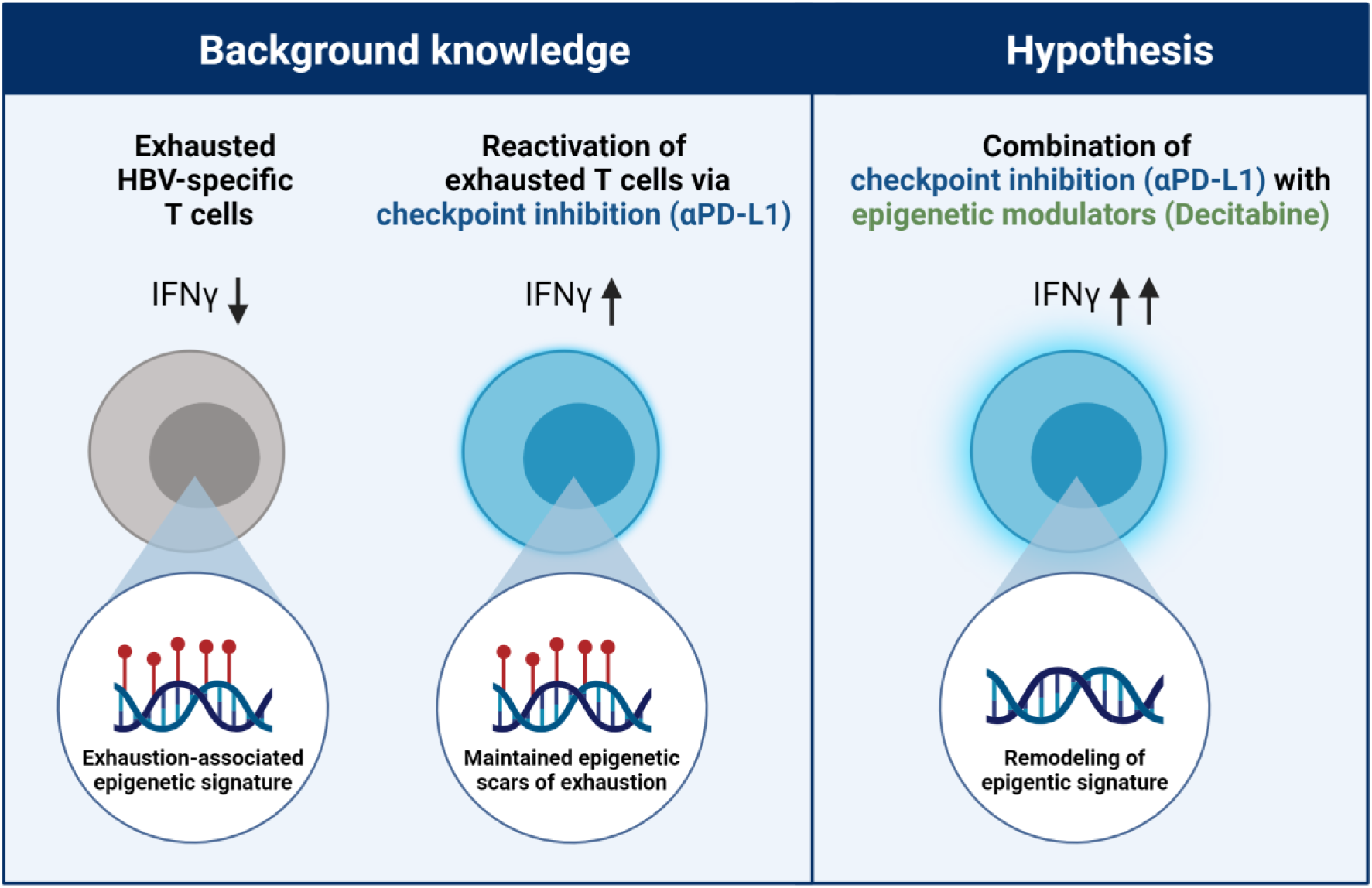

**Highlights:**

- Remodeling epigenetic signatures with a DNA methyltransferase inhibitor enhances the effectiveness of immune checkpoint inhibition in restoring HBV-specific T cell responses.
- Responsiveness is associated with specific IFNγ DNA methylation patterns and plasma levels.
- Epigenetic remodeling had distinct effects on two CD8 T cell epitopes, with more pronounced effects on HBV-pol_455_-specific CD8^+^ T cell responses.

## Introduction

Infection with the hepatitis B virus (HBV) is a relevant public health challenge, with more than 250 million individuals worldwide living with chronic infection [1]. Current antiviral therapies, e.g., with nucelos(t)ide analogues, are effective in viral suppression but rarely achieve the ultimate goal of functional cure defined as loss of HBV surface antigen (HBsAg) [2]. The persistent presence of antigens in chronic diseases leads to T cell exhaustion, marked by reduced cytokine production (e.g., IFNγ), elevated levels of inhibitory receptors like PD-1, and alterations in transcriptional and epigenetic signatures [3–6].

One approach to restore exhausted HBV-specific T cells is to employ immune checkpoint inhibitor (ICI) therapy to block the PD-1 interaction, such as with αPD-1 or αPD-L1 antibodies [7–9]. This approach represents a major breakthrough in the field of immunotherapy and is widely used for the treatment of various cancers [10], including hepatocellular carcinoma (HCC) due to chronic HBV infection [11]. ICI therapy has also been prospectively evaluated in patients without HCC and chronic HBV infection to achieve a functional cure, albeit with limited efficacy to date [12].

Epigenetic signatures linked to T cell exhaustion may diminish the effectiveness of ICI therapy. These persistent epigenetic imprints are evident in various chronic viral infections, including lymphocytic choriomeningitis virus (LCMV) and chronic hepatitis C virus (HCV), in both mice and humans. Even though LCMV and HCV infections can be cured (unlike chronic HBV), the epigenetic scars often persist long-term after antigen removal, continuing to affect the immune system [13–15]. Moreover, these epigenetic imprints are not reversed by αPD- L1 checkpoint inhibition [5,16].

Subsequently, targeting epigenetic mechanisms might be one approach to improve ICI efficacy [17]. The most prominent type of epigenetic modification is DNA methylation, which is catalyzed by DNA methyltransferases (DNMT). Recently, there has been evidence that *de novo* DNA methylation is a crucial mechanism for T cell exhaustion development [18] and is therefore a reasonable target for new therapies. DNMT inhibitors can block *de novo* DNA methylation leading to demethylation. In cancer, clinical trials combining DNMT inhibitors with checkpoint inhibitors are ongoing and have already resulted in improved clinical response rates [19,20]. Decitabine (DAC) is a potent inhibitor of DNMTs and subsequent DNA methylation. DAC is already approved for the treatment of hematological malignancies such as myelodysplastic syndrome and acute myeloid leukemia [21]. Recent studies have shown that combining DAC pretreatment with the ICI αPD-L1 can remodel exhaustion-associated epigenetic patterns and reactivate impaired T cells in the chronic LCMV infection mouse model [18]. In chronic HBV infection, the potential of epigenetic remodeling to improve the efficacy of ICI to restore T cell responses has not yet been investigated and demonstrated. The aim of this study was therefore to investigate whether the combination of the DNMT inhibitor DAC with the ICI αPD-L1 can alter HBV-specific T cell responses in chronic HBV infection. The goal was to underscore the role of epigenetic signatures in HBV-specific T cells and explore a novel strategy for improving their response.

## Patients and methods

### Patients and sample preparation

Three cohorts of patients with chronic HBV infection (CHB) were assigned for this study (Table 1). Clinical parameters were extracted from the Enterprise Clinical Research Data Warehouse of the Hannover Medical School. Viral HBsAg and HBcrAg levels were measured as previously described [22], using the Abbott ARCHITECT and Lumipulse G assays, respectively.

**Table 1:**
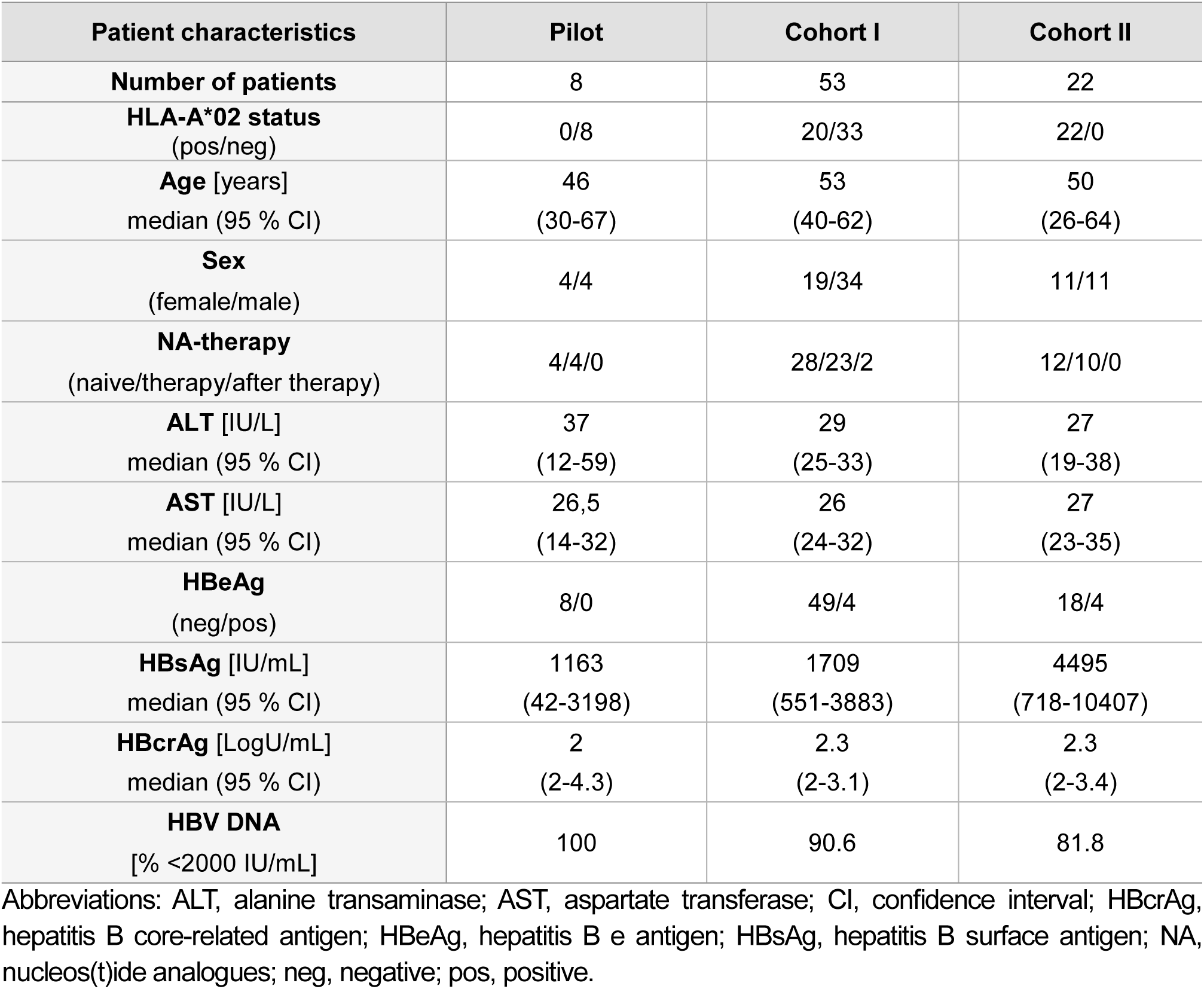
Study cohorts of patients with chronic HBV infection.

| Patient characteristics | Pilot | Cohort I | Cohort II |
| --- | --- | --- | --- |
| **Number of patients** | 8 | 53 | 22 |
| **HLA-A\*02 status**<br>(pos/neg) | 0/8 | 20/33 | 22/0 |
| **Age [years]**<br>median (95 % CI) | 46<br>(30-67) | 53<br>(40-62) | 50<br>(26-64) |
| **Sex**<br>(female/male) | 4/4 | 19/34 | 11/11 |
| **NA-therapy**<br>(naive/therapy/after therapy) | 4/4/0 | 28/23/2 | 12/10/0 |
| **ALT [IU/L]**<br>median (95 % CI) | 37<br>(12-59) | 29<br>(25-33) | 27<br>(19-38) |
| **AST [IU/L]**<br>median (95 % CI) | 26,5<br>(14-32) | 26<br>(24-32) | 27<br>(23-35) |
| **HBeAg**<br>(neg/pos) | 8/0 | 49/4 | 18/4 |
| **HBsAg [IU/mL]**<br>median (95 % CI) | 1163<br>(42-3198) | 1709<br>(551-3883) | 4495<br>(718-10407) |
| **HBcrAg [LogU/mL]**<br>median (95 % CI) | 2<br>(2-4.3) | 2.3<br>(2-3.1) | 2.3<br>(2-3.4) |
| **HBV DNA**<br>[% <2000 IU/mL] | 100 | 90.6 | 81.8 |

Abbreviations: ALT, alanine transaminase; AST, aspartate transferase; CI, confidence interval; HBcrAg, hepatitis B core-related antigen; HBeAg, hepatitis B e antigen; HBsAg, hepatitis B surface antigen; NA, nucleos(t)ide analogues; neg, negative; pos, positive.

Plasma and blood samples were collected from CHB patients. PBMCs were isolated and cryopreserved for later use as described elsewhere [23]. *Ex vivo* plasma IFNγ level was determined by Olink proteomics analysis (Uppsala, Sweden).

All experiments were carried out in accordance with the principles espoused in the Declaration of Helsinki. The local ethics committee of Hannover Medical School ensured this project. Written informed consent was obtained from all participants.

### HBV peptides and DNA methyltransferase inhibitor decitabine

All peptides were synthesized by ProImmune (Oxford, UK). The HBV core overlapping peptide pool (OLP) consists of 15 amino acid long peptides that overlap by 10 amino acids spanning the entire HBV core antigen (41 peptides) of HBV Genotype D ayw (U95551.1). After solubilization in 100% DMSO, the peptide pool was further diluted 1:6 in HBSS (Gibco, CA, USA) to reach a final concentration of 1.25 µg/mL per peptide. HLA-A*02-restricted HBV-specific peptides, core_18-27_ (core_18_, FLPSDFFPSV) and polymerase_455-463_ (pol_455_, GLSRYVARL) were solubilized in 100% DMSO and used in a final concentration of 10 µg/mL.

Decitabine (DAC, solubilized in 100% DMSO, Selleckchem, Planegg, Germany) was diluted in 100% DMSO to reach final concentrations of 0.1, 0.3 and 1 µM.

### Modulation of HBV-specific T cell response after in vitro stimulation with HBV-specific peptides

HBV-specific T cell responses were analyzed as previously described [24] (for detailed antibodies and procedure, see online supplementary material). Briefly, PBMCs were stimulated for 10 days with HBV core OLP or HLA-A*02-restricted peptides (core_18_ and pol_455_). Different combinations of DAC with or without αPD-L1 were used as treatments. IL2- containing medium was provided at days 3 and 7 post-stimulation. After 10 days, cells were restimulated with the respective peptides with or without αPD-L1 treatment. Intracellular staining was performed after fixation and data were acquired on a BD LSRFortessa flow cytometer. The T cell responses were calculated by subtracting the medium background control.

In cohort 1, patients were categorized as DAC-responders or non-responders based on their IFNγ CD4^+^ T cell response after 10 days of culture with different treatments. DAC- responders exhibited a higher IFNγ CD4^+^ T cell response with the combination of DAC and αPD-L1 compared to αPD-L1 monotherapy or core OLP alone. Specifically, the IFNγ response was calculated as follows: *x*_core_, *x*_*αPD−L*1_, and *x*_*DAC*_ represent the percentage of IFNγ^+^ cells in CD4^+^ T cells for core OLP, αPD-L1 monotherapy, and the DAC/αPD-L1 combination, respectively, with the percentage of IFNγ^+^ cells in CD4^+^ T cells in the medium control subtracted. We then calculated the IFNγ response for each condition. The values ^are as follows: Δ^αPD−L1 ^= *x*^αPD−L1 ^− *x*^core^, Δ^DAC1 ^= *x*^DAC ^− *x*^core^, and Δ^DAC2 ^=*x*^DAC ^− *x*^αPD−L1. Finally, the patients are assigned to each group based on the following criteria.

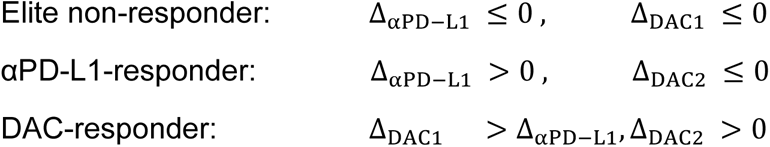

Elite non-responders and αPD-L1–responders are further defined as non-responders.

### DNA methylation analysis

DNA was isolated from PBMCs by using Monarch Genomic DNA Purification Kit T3010L (New England Biolabs, Ipswich, MA, USA) and methylation was analyzed at Human Genomics Facility of Erasmus MC (The Netherlands, Rotterdam) using the Infinium MethylationEPIC BeadChip array.

Methylation data was processed in R software (v2022.07.0) as previously described [25]. Briefly, DNA methylation values calculated from the original IDAT files using the minfi [26] package in R (v.4.2.0). Probes with missing methylation values (>10%), those on sex chromosomes, those containing single nucleotide polymorphisms (MAF>5%), and probes mapping to multiple loci were excluded. In total, 73,308 unique probes failed and were removed from the dataset. After quality control and “dasen” normalization from the “watermelon ” package [27], 35 samples and 792,930 probes remained for downstream analysis.

### Differential methylation analysis

Genomic position of each methylated sites, i.e. CpGs, were annotated using “IlluminaHumanMethylationEPICanno.ilm10b4.hg19”. The location of genes and their promoter regions based on human reference genome GRCh37/hg19 were obtained from UCSC Genome Browser (https://genome.ucsc.edu/). Mean methylation level of the respective genes were plotted. Methylation trend of DAC-responders was calculated by subtracting the methylation level of non-responders.

### Statistical analysis

Statistical analyses were performed using GraphPad Prism 8.4.2 (USA, CA) and R software. Statistical significance was tested depending on the type of data as indicated in the figure legends. Non-parametric data: Wilcoxon matched-pairs signed rank test, Mann-Whitney test, Friedman test, and Spearman correlation. Parametric data: unpaired t-test, mixed- effects model, and Pearson correlation. All tests were carried out as 2-tailed tests. Significant p-values were considered as follows: *p<0.05; **p<0.01; ***p<0.001; ****p<0.0001.

The effect size Cohen’s d was calculated between two groups by using the differences between the means divided by the pooled standard deviation. The effect was defined as small (*d*=0.2-0.5), medium (*d*=0.5-0.8), and large (*d*>0.8).

The fold change of the immune responses was calculated after the addition of 0.01 to all values to include peptide stimulation non-responders.

## Results

### Cytotoxicity and immunomodulatory effects of different decitabine (DAC) concentrations *in vitro*

First, we assessed the treatment conditions to study the immunomodulatory effect of the combination of DAC with αPD-L1. PBMCs from 8 CHB patients were treated with various DAC concentrations (Table 1 Pilot cohort, Figure 1A). Flow cytometry analysis showed decreased live lymphocyte frequency with DAC concentrations >0.3 µM, while 0.1 µM DAC had no detectable cytotoxicity (Figure 1B-C). DAC monotreatment showed no significant impact on CD4^+^ and CD8^+^ T cell IFNγ immune response (Figure 1D). Therefore, experiments proceeded with 0.1 µM DAC in combination with αPD-L1. The comparison of simultaneous DAC and αPD-L1 administration versus DAC-pretreatment revealed that the pretreatment approach significantly augmented IFNγ response and the multifunctionality, as demonstrated by CD4^+^ T cells in the majority of the 36 tested CHB patients (Figure 1E-G).

**Figure 1:**
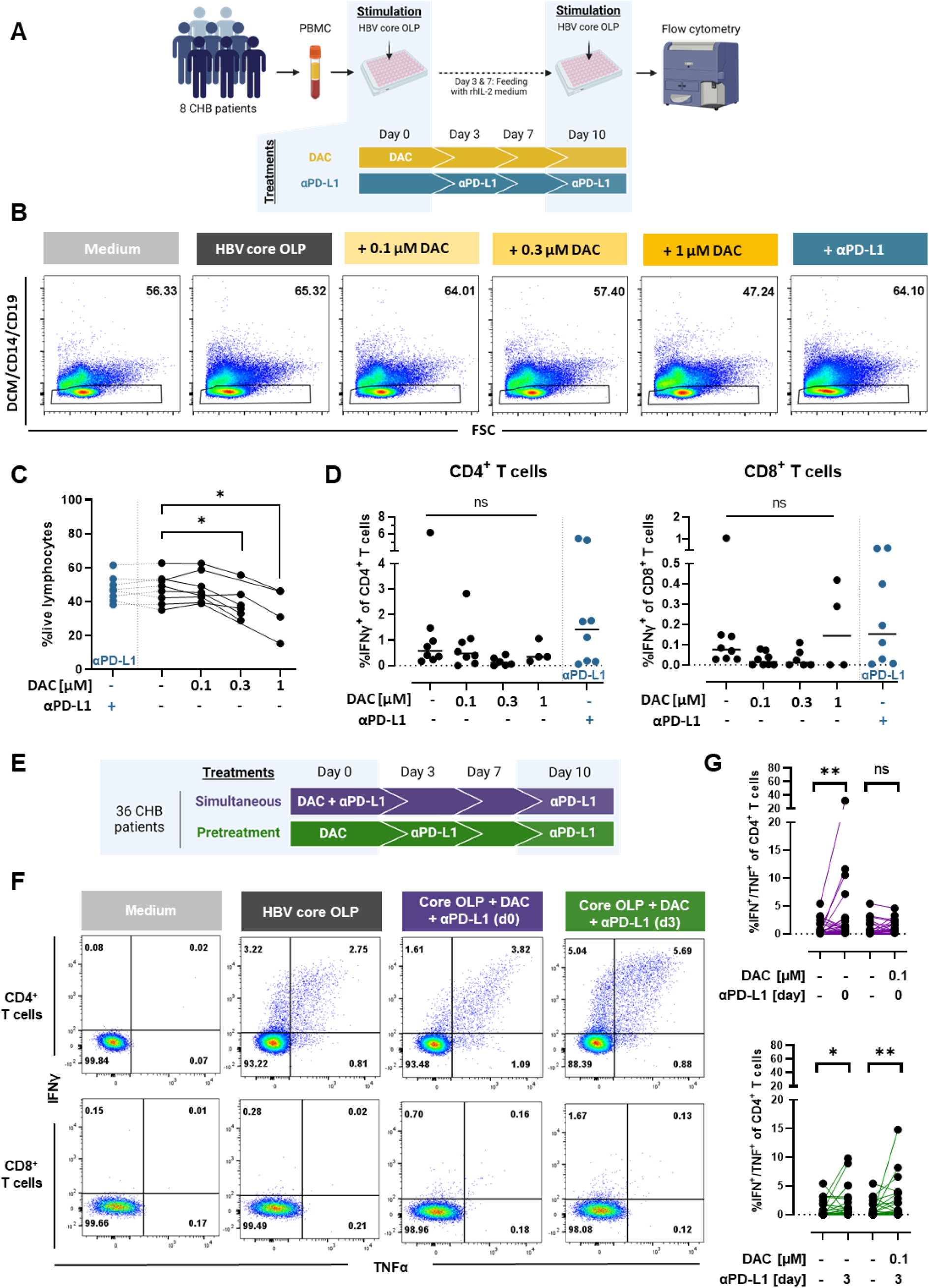
Cytotoxic potential of decitabine (DAC) as monotreatment and combinatory immunomodulatory effect with αPD-L1 *in vitro*. **(A)** Overview of 10-day culture with HBV core OLP stimulation and different DAC concentrations (n=8). **(B)** Representative flow cytometry plots for live lymphocytes (DCM^-^/CD14^-^/CD19^-^) in each condition **(C)** Frequency of live lymphocytes (DCM^-^/CD14^-^/CD19^-^) within total PBMCs in response to increasing concentrations of DAC compared to αPD-L1 monotreatment (depicted in blue). **(D)** IFNγ response of CD4^+^ and CD8^+^ T cells following exposure to varying doses of DAC or αPD-L1. **(E)** Overview of 10-day culture with HBV core OLP stimulation and DAC as pretreatment or simultaneous addition with αPD-L1 (n=36). **(F)** Representative flow cytometry plots of one patient depicting IFNγ/TNFα response of CD4^+^ and CD8^+^ T cells after different treatment approaches. **(G)** Quantification of IFNγ/TNFα response of CD4^+^ T cells. Statistical significance was determined using mixed-effects analysis with Geissner-Greenhouse correction and Dunnett’s multiple comparison test (C-D) and Wilcoxon matched-pairs signed rank test (G) (ns: not significant; p*<0.05; p**<0.01).

### Immunomodulatory effect of DAC and αPD-L1 on HBV core OLP-stimulated immune cells

Therefore, a low dose DAC-pretreatment approach was pursued in a total of 53 CHB patients (Table 1 cohort I, Figure 2A). Both CD4^+^ and CD8^+^ T cell subsets exhibited correlated immune responses to the different treatments; however, CD4^+^ IFNγ responses were more pronounced than CD8^+^ T cell responses (Figure 2B-E). This discrepancy may be attributed to the more restricted binding cleft of MHC I and the length of the HBV core OLP (15-mers) hindering optimal recognition and activation of CD8^+^ T cells. Consequently, CD8^+^ T cell responses were investigated separately with HLA-A*02-restricted HBV epitopes in the following section of this study. The investigation of core OLP-stimulated samples focused on the CD4^+^ T cell responses. Overall, the CD4^+^ IFNγ response to the different treatments was quite heterogeneous. Nine patients responded strongly to the DAC/αPD-L1 combination, with CD4^+^ IFNγ responses increasing significantly compared to HBV core OLP stimulation alone (Figure 2F). Moreover, compared to αPD-L1 monotreatment, the CD4^+^ immune response of 12 out of 53 patients was enhanced by a factor of 2-100 (Figure 2G). Overall, the effect size of the combination treatment was greater compared to HBV core OLP (*d*=0.26) and αPD-L1 monotherapy (*d*=0.15). Approximately 32% of patients met the criteria for DAC responsiveness, characterized by a higher frequency of DAC-mediated IFNγ^+^ CD4^+^ response compared to peptide stimulation and αPD-L1 monotherapy. Of note, the response pattern to DAC/αPD-L1 was consistent in an independent experiment with a different investigator using a refrozen sample (Supplementary Figure 1). In summary, the DAC/αPD-L1 combination led to a heterogeneous patient-dependent response and has the potential to improve the CD4^+^ IFNγ response in a subset of patients.

**Figure 2:**
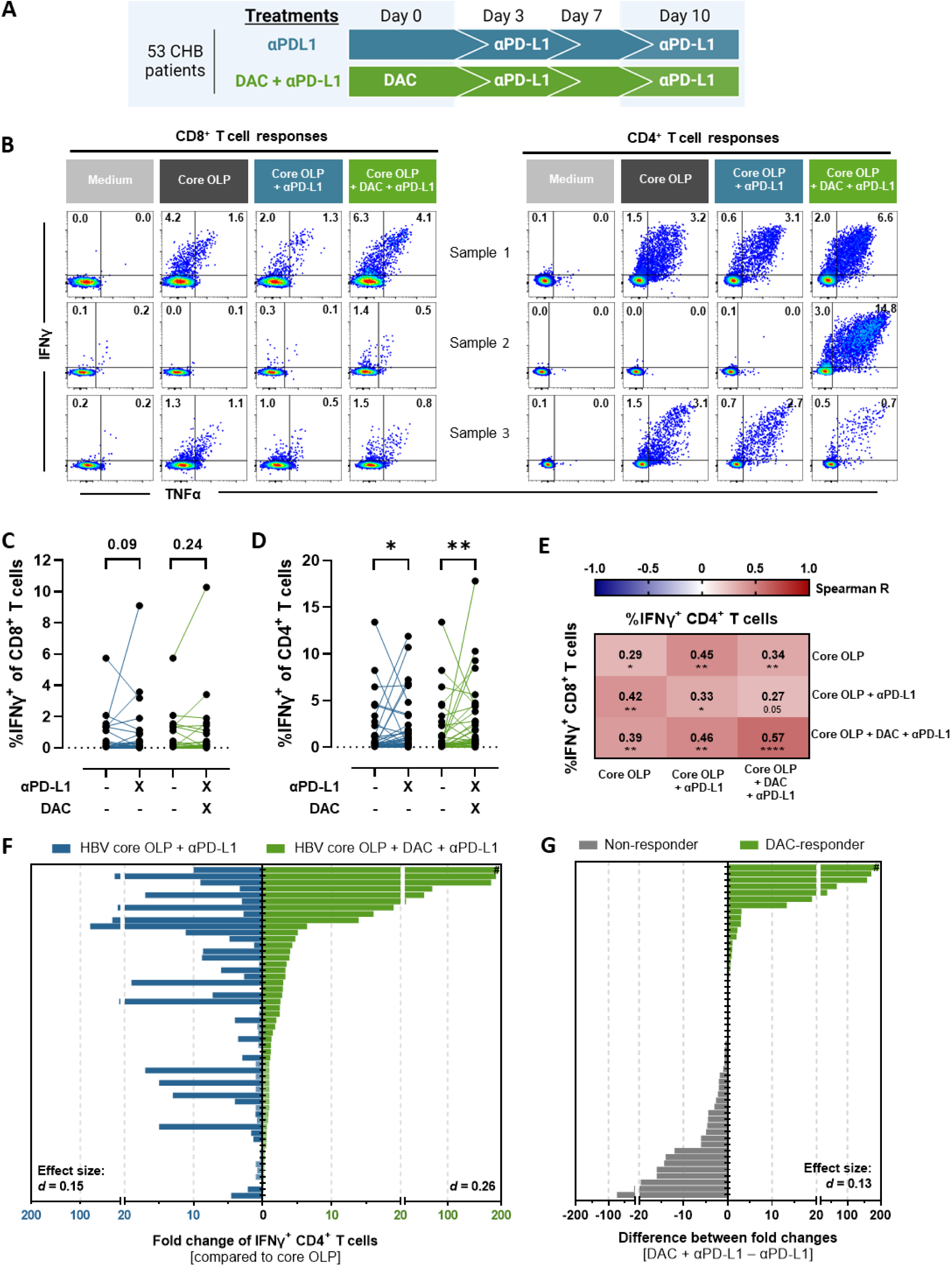
Heterogeneous immunomodulatory effect of DAC in combination with αPD-L1 on HBV core OLP-stimulated PBMCs. **(A)** Treatment overview of the 10-day culture with HBV core OLP stimulation. **(B)** Flow cytometry plots of CD8^+^ and CD4^+^ IFNγ/TNFα response in DAC-responders (sample 1-2) and non-responder (sample 3). **(C-D)** CD8^+^ (C) and CD4^+^ (D) T cell IFNγ responses after core OLP 10-day culture of 53 patients. **(E)** Spearman correlation matrix of CD8^+^ and CD4^+^ IFNγ^+^ T cell responses. **(F)** Fold change of IFNγ^+^ CD4^+^ T cells compared to HBV core OLP stimulated control upon αPD-L1 treatment (blue) or combination of DAC and αPD-L1 (green). Fold change was determined after adding 0.01 to all values to include HBV core OLP-non-responders. **(G)** Difference between fold changes of DAC/αPD-L1 combination and αPD-L1. Statistical significance was determined using Wilcoxon matched-pairs signed rank test (p*<0.05; **p<0.01) and Cohen’s *d* effect size. #: Fold change/Difference >1500.

### Epitope-specific impact of DAC and αPD-L1 combination on HBV-specific CD8^+^ T cell responses

Next, we explored the CD8^+^ T cell response to two model epitopes, known to differ in their exhaustion levels [28]. PBMCs from HBV patients (Table 1 cohort II) were stimulated with HLA-A*02-restricted peptides, core_18_ and pol_455_ (Figure 3A). Distinct epitope-specific responses were observed in IFNγ production upon stimulation with corresponding peptides (Figure 3B-C). Core_18_-stimulated CD8^+^ T cells produced more IFNγ than pol_455_-specific CD8^+^ T cells. The addition of αPD-L1 positively impacted the IFNγ response of both core_18_- and pol_455_-specific CD8^+^ T cells, with a more homogeneous enhancing effect observed in the latter, corroborating our previous findings [24]. Furthermore, the novel strategy involving DAC-pretreatment followed by αPD-L1 treatment enhanced the IFNγ response of both core_18_- and pol_455_-specific CD8^+^ T cell populations in 7 and 10 patients, respectively. Particularly, the combination of DAC and αPD-L1 increased the IFNγ production in pol_455_- specific CD8^+^ T cells in nearly half of the tested patients (10 DAC-responders vs. 12 non-responders). Compared to αPD-L1 monotreatment, the DAC/αPD-L1 combination showed a significant effect in the pol_455_-specific immune response (effect size: core_18_ *d*=0.1, pol_455_ *d*=0.47).

**Figure 3:**
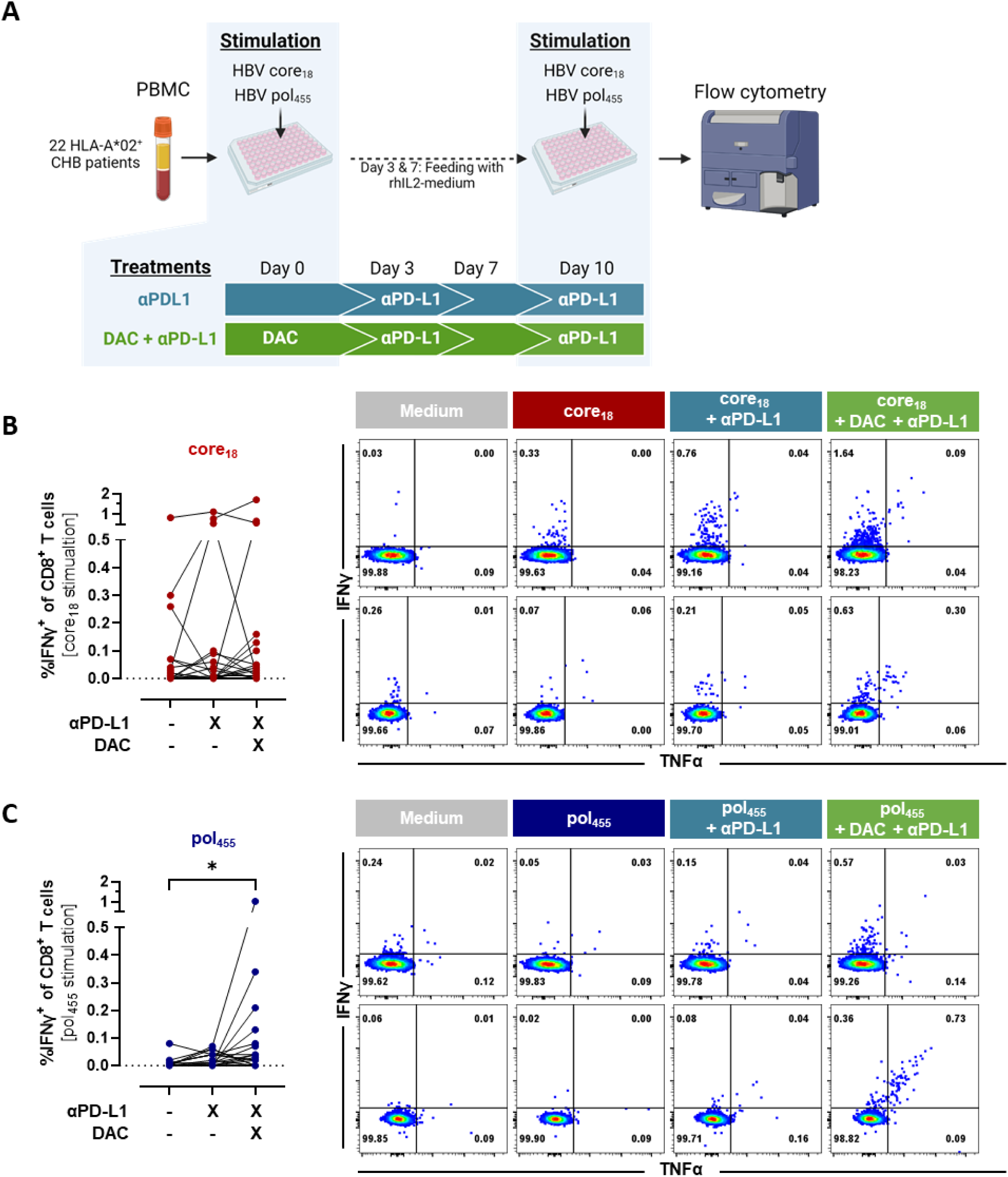
Augmented HBV-specific CD8^+^ T cell responses with combined DAC and αPD-L1 treatment. (**A**) Schematic overview of experimental setup. (**B-C**) IFNγ response of CD8^+^ T cells and representative flow cytometry plots of two DAC-responders upon core18 (**B**) or pol455 (**C**) stimulation. Statistical significance was determined using Friedman test with Dunn’s multiple comparison (p*<0.05).

In summary, responsiveness to combined DAC/αPD-L1 treatment differs between HBV epitope-specific CD8^+^ T cell populations, with the combination showing particular efficacy in boosting pol_455_-specific CD8^+^ T cell responses.

### Characteristics of CD4^+^ T cell DAC-responders and non-responders

The heterogeneous immune responses to the DAC/αPD-L1 combination underscore the need to identify responders and non-responders for effective immunotherapy. To pinpoint predictive markers for this approach, we compared DAC-responders and non-responders from cohort I, which had a larger sample size (Table 1). DAC-responders were characterized by a higher IFNγ^+^ CD4^+^ T cell response following DAC/αPD-L1 treatment compared to peptide stimulation and αPD-L1 monotherapy, while non-responders showed no improvement in IFNγ^+^ CD4^+^ T cell response with the combination treatment. As defined, DAC-responders (n=17) exhibited robust IFNγ^+^ CD4^+^ responses following combined DAC and αPD-L1 treatment (Supplementary Figure 2A), significantly higher than non-responders (n=36, p=0.005). Conversely, non-responders responded significantly to αPD-L1 treatment, with DAC-pretreatment apparently exerting a counteractive effect on the immune response. While responses to initial HBV core stimulation or αPD-L1 treatment were slightly higher in the non-responder group, this difference was not statistically significant between the two groups.

Since age, medication and virological markers might influence the immune response [24,29], we first compared clinical characteristics between DAC-responders and non-responders. However, none of the investigated clinical parameters including HBsAg and HBcrAg levels, differed significantly between DAC-responders and non-responders (Table 2) nor correlated significantly with the IFNγ response upon the DAC/αPD-L1 combination treatment. Nevertheless, we noticed that the response to αPD-L1 monotreatment decreased with increasing HBcrAg levels (known from our previous study [24]), whereas the response to DAC/αPD-L1 appeared to be independent of HBcrAg level (Figure 4).

**Figure 4:**
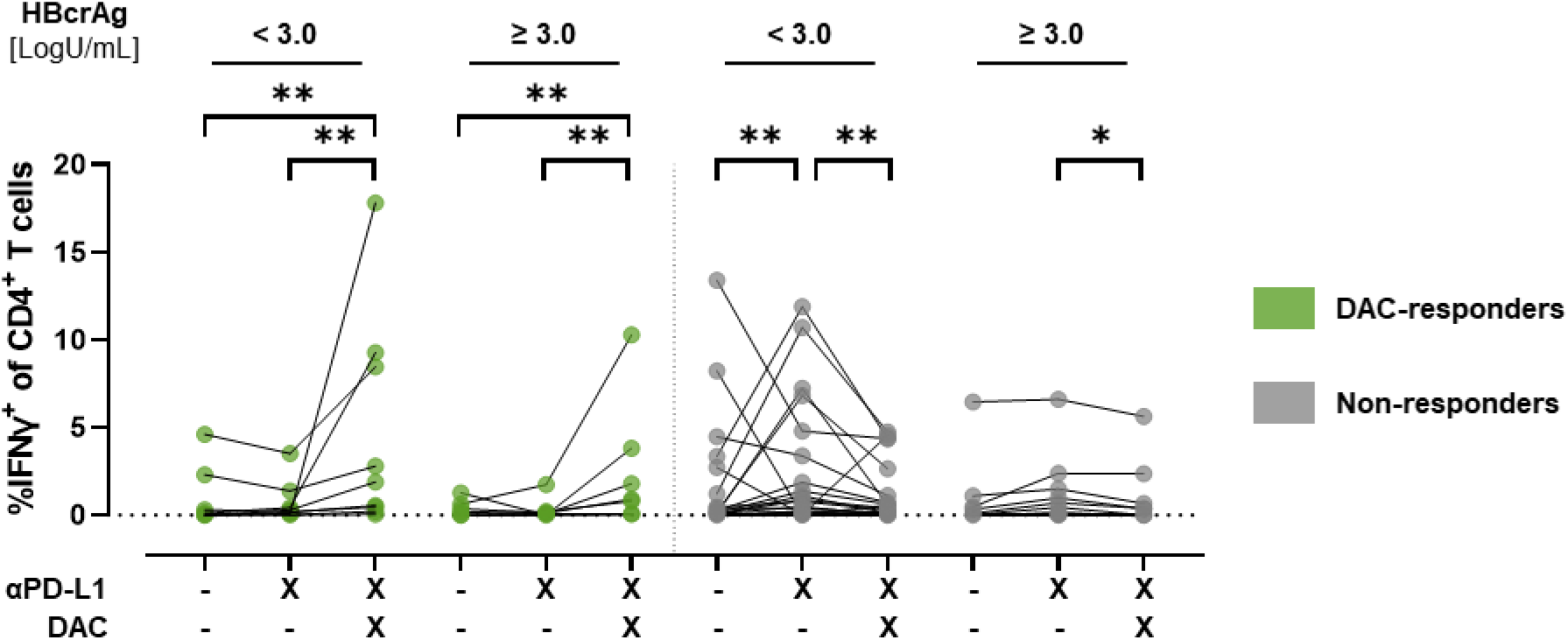
Role of HBV core-related antigen (HBcrAg) in responsiveness. CD4^+^ IFNγ response after HBV core OLP 10-day culture (from Figure 2D) divided based on DAC-responsiveness (DAC-responders in green; Non-responders in grey) and HBcrAg level. Statistical significance was determined using Friedman test with Dunn’s multiple comparison (p*<0.05; p** <0.01).

**Table 2:** Clinical characteristics of DAC-responders and non-responders.

|  | DAC-responder | Non-responder |
| --- | --- | --- |
| **Number of patients** | 17 | 36 |
| **Age** [years]<br>median (95 % CI) | 53<br>(37-67) | 54.5<br>(40-63) |
| **Sex**<br>(female/male) | 7/10 | 12/24 |
| **Therapy**<br>(naive/therapy/a.t.) | 10/5/2 | 18/18/0 |
| **ALT** [IU/L]<br>median (95 % CI) | 29<br>(24-39) | 28.5<br>(22-32) |
| **AST** [IU/L]<br>median (95 % CI) | 29<br>(24-35) | 26<br>(23-32) |
| **HBeAg**<br>(neg/pos) | 15/2 | 34/2 |
| **HBsAg** [IU/mL]<br>median (95 % CI) | 1939<br>(176-10407) | 1571<br>(718-3883) |
| **HBcrAg** [LogU/mL]<br>median (95 % CI) | 2.5<br>(2-3.5) | 2<br>(2-3.1) |
| **HBV DNA**<br>[% <2000 IU/mL] | 94 | 92 |

Abbreviations: ALT, alanine transaminase; AST, aspartate transferase; a.t, after therapy; CI, confidence interval; DAC, decitabine; HBcrAg, hepatitis B core-related antigen; HBeAg, hepatitis B e antigen; HBsAg, hepatitis B surface antigen; neg, negative; pos, positive.

Furthermore, the proportion of patients receiving antiviral therapy with nucleos(t)ide analogues (NA) was lower in the DAC-responder group (30%) compared to the non-responder group (49%). Additionally, analysis of IFNγ response following combined DAC/αPD-L1 treatment showed that patients receiving NA-therapy had lower responses compared to therapy-naïve individuals (p=0.03). Although not reaching statistical significance, immune responses to HBV core OLP or αPD-L1 monotreatment were also lower in patients undergoing NA-therapy (p=0.07 and p=0.09, respectively). Therefore, a bias related to the reason for NA therapy may contribute to the observed variation in immune responses. However, some patients on NA-therapy were still among the highest DAC- responders, suggesting that NA-therapy alone is not a dominant factor.

### *Ex vivo* epigenetic characteristics of treatment-responsiveness to DAC/αPD-L1

Given DAC’s role as an epigenetic modifying drug, we hypothesized that the heterogeneity of the immune responses may be explained by epigenetic differences of the immune cells between DAC-responders and non-responders. The low frequency of HBV antigen-specific T cells hinders direct investigation of epigenetic modification in the PBMC amount that was available from the patients. Therefore, we opted to examine the epigenetic profiles in bulk PBMC instead. First, we investigated the methylation of genes associated with epigenetic modifications (e.g. *DNMT*s*, HDAC*s*, TET*s), immune cell functionality (e.g. *IFNG* (encoding for IFNγ), *PDCD1* (encoding for PD-1), *CD274* (encoding for PD-L1)), and antigen presentation machinery (e.g. *TAP*s, *PSMB*s). DAC-responders (n=11) exhibited an overall more hypermethylated state than non-responders (n=24), particularly in genes associated with immune cell functionality and antigen presentation machinery (Figure 5A). Of note, the methylation trend in the DAC-responders positively correlated with the initial methylation level of the selected genes (Figure 5B). This suggests that hypermethylated genes were more likely to exhibit varying levels of methylation between the groups compared to the more hypomethylated genes of the DAC-responders.

**Figure 5:**
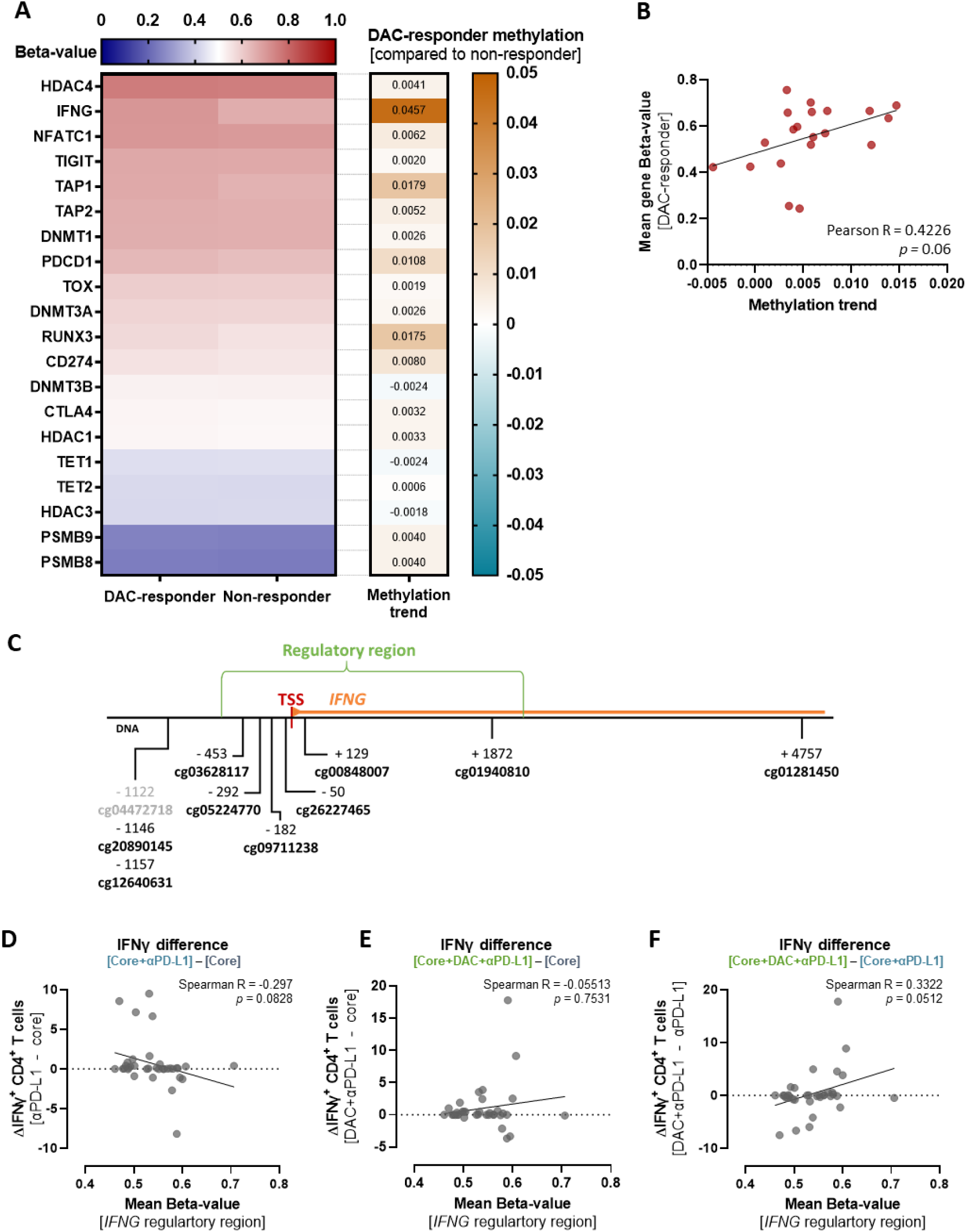
Methylation analysis of *IFNG* in DAC-responders compared to non-responders. (**A**) Mean gene methylation (left panel) and methylation trend in DAC-responders compared to non-responders (right panel). (**B**) Pearson correlation of mean gene methylation of DAC-responders and detected methylation trend (**p<0.01). (**C**) Schematic representation of the genomic location of CpG sites associated with *IFNG* in relation to the transcription start site (TSS) and promotor-associated regulatory region (GH12J068159). Black marked CpG sites indicate higher methylation in DAC-responders. Numbers indicate distance to TSS in base pairs. (**D-F**) Spearman correlation analysis depicting the relationship between the mean methylation levels of *IFNG* regulatory region-associated CpG sites and the difference in IFNγ expression in CD4^+^ T cells between different stimulations (n = 35).

*IFNG* methylation displayed the most significant methylation difference among the selected genes. Thus, we further investigated the methylation level of *IFNG*-associated CpG-sites in more detail (Figure 5C). Almost all *IFNG*-associated CpG-sites (despite cg04472718) were more methylated in the DAC-responder group. The majority of CpG sites were located in a promotor-associated regulatory region (GH12J068159). The mean methylation of the regulatory region did not correlate with the detected total IFNγ response of CD4^+^ T cell after 10-day stimulation cultures (Spearman R=0.23; p=0.19). Instead, the *ex vivo* methylation level of the regulatory region appears to be associated with the difference in IFNγ response between the different treatments used, i.e. the treatment effect (Figure 5D-F). Although not statistically significant, a lower level of methylation of the *IFNG* regulatory region was associated with a greater difference in response to αPD-L1 compared to the core OLP response alone (Figure 5D), while a higher level of methylation was associated with a more pronounced difference in DAC/αPD-L1 response compared to αPD-L1 (Figure 5F). This suggests that methylation of the *IFNG* regulatory region essentially influences the response to αPD-L1 checkpoint inhibition. To support this hypothesis of an association of *IFNG* methylation to αPD-L1 responsiveness, we further divided the gene-specific methylation of the non-responder group into αPD-L1-responders, initially responsive to αPD-L1 but showing reduced response with DAC, and elite non-responders, unresponsive to both treatments (Supplementary Figure 2B). Methylation levels of key genes *IFNG* and *PDCD1* were compared among these groups (Supplementary Figure 2C). Intriguingly, αPD-L1- responders showed significant lower *IFNG* methylation compared to DAC-responders and elite non-responders (p=0.018, p=0.018, respectively). No significant differences were observed between DAC-responders and elite non-responders in *IFNG* methylation. This *ex vivo* methylation signature of the subgroups reflected the IFNγ^+^ CD4^+^ T cell immune response and responsiveness to checkpoint inhibition observed after 10-day cultures (Supplementary Figure 2B). Conversely, *PDCD1* expression was similar in αPD-L1- responders and elite non-responders, distinguishing both from DAC-responders (p=0.0001, p=0.0001, respectively). DAC-responders exhibited higher methylation levels in both selected genes.

These epigenetic analyses highlight a unique epigenetic profile in DAC-responders and suggest that αPD-L1-responsiveness might be linked to less methylated *IFNG* gene.

### *Ex vivo* plasma IFNγ level as marker for DAC/αPD-L1 response

Given that epigenetic imprints are associated with distinct transcriptional activity, we additionally investigated the *ex vivo* IFNγ plasma level of the respective patients. Out of 35 patients with methylation data available, 29 passed the Olink analysis quality control. In accordance with the methylation data (higher methylation of the *IFNG* genes), DAC- responders exhibited significantly lower IFNγ plasma levels compared to non-responders (Figure 6A). The IFNγ plasma level correlated not only with the *ex vivo* IFNγ methylation (Spearman R = -0.3; p = 0.05), but also with the *in vitro* CD4^+^ T cell immune response only after combined DAC/αPD-L1 treatment (Figure 6B-C). It can be concluded that the CD4^+^ T cell immune response to DAC and αPD-L1 is higher if the *ex vivo* IFNγ plasma level is low and *IFNG* is more hypermethylated.

**Figure 6:**
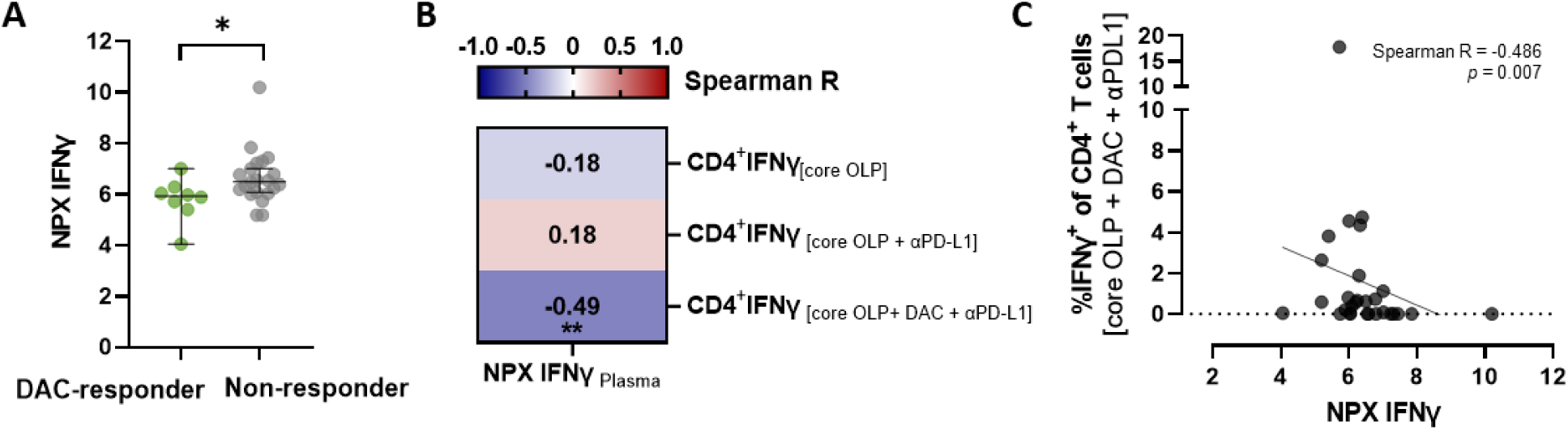
Plasma IFNγ level as marker for DAC/αPD-L1 response. **(A)** IFNγ plasma level of DAC- responders (n=8) and non-responders (n=21). **(B)** Spearman correlation of *ex vivo* IFNγ plasma level and CD4^+^ T cell IFNγ responses (n=29) after 10-day culture with the respective treatments. **(C)** Correlation of *ex vivo* IFNγ plasma level and frequency of IFNγ^+^ CD4^+^ T cells after 10-day culture with HBV core OLP stimulation and combined DAC/αPD-L1 treatment. Statistical significance was determined using the Mann- Whitney test (A) and Spearman correlation (B-C) (**p<0.01).

## Discussion

In recent years, research on epigenetic signatures and alterations has advanced, providing a basis for potential therapeutic breakthroughs that have so far been mainly apparent in cancer treatment [17]. However, exhausted immune responses are also closely associated with epigenetic imprints in chronic infectious diseases [18]. Therefore, one potential approach to restore exhausted T cell responses involves targeting epigenetics, such as employing DNA methyltransferase inhibitors like decitabine (DAC), in synergy with checkpoint inhibitors [18]. Despite its promising results in various scenarios, the application and potential of this strategy in chronic HBV infection has not been explored and has so far not informed current therapeutic considerations regarding HBV cure strategies.

This study demonstrates that the combination of the DNA methyltransferase inhibitor DAC and the ICI αPD-L1 enhances HBV-specific CD4^+^ and CD8^+^ T cell responses *in vitro*, while DAC monotherapy alone could not restore these responses, consistent with findings in the LCMV infection mouse model [18]. Similar to previous studies [30,31], we observed a dose- dependent reduction in live lymphocytes, likely due to DAC’s cytotoxicity. At 0.1 µM, DAC has demonstrated hypomethylating effects with maximal demethylation occurring after 72 hours [32], which could explain the improved effect of DAC-pretreatment compared to simultaneous treatment. This time-consuming demethylation step of DAC is necessary to improve the response to αPD-L1 and is therefore not detectable in *ex vivo* functional assays.

Despite promising T cell responses in some patients, the response to the combination was heterogeneous. A subset of patients exhibited significantly enhanced immune responses compared to αPD-L1 alone, while others showed no response or a decline in their already robust immune response. This variability aligns with previous reports of ICI use *in vitro* [24] or in patients [12], highlighting the diversity of immune responses in chronic HBV infection [8]. The main factor that distinguished responder to DAC/αPD-L1 treatment from non- responder was the epigenetic signature, in particular in the *IFNG* gene. DAC-responders exhibited a constrained response to αPD-L1, potentially due to hypermethylated regions crucial for checkpoint inhibition. DAC, as a demethylating agent, can counteract these hypermethylated imprints, thereby enhancing finally the observed T cell activation. However, a comprehensive understanding of the treatment’s exact mechanism requires further exploration at the epigenetic level. Validating the epigenetic changes induced by the DAC/αPD-L1 combination will not only enhance mechanistic insights but also help identify more precise targets for intervention.

Notably, previous studies indicated that patients with lower antigen levels (e.g., HBsAg <100 IU/mL or HBcrAg <3.87 LogU/mL) responded better to ICI *in vitro* [24], and those with lower HBsAg levels also responded better to ICI therapy *in vivo* [33]. Indeed, HBV peptide stimulation and αPD-L1 monotreatment were mostly effective only in patients with low HBcrAg levels, consistent with previous results [24]. Importantly, we found no differences in HBsAg or HBcrAg levels between responders and non-responders to the DAC/αPD-L1 combination in our study, suggesting that this treatment can overcome the greater immune exhaustion in patients with higher HBV antigen levels. Conversely, the DAC/αPD-L1 combination showed strong effects even in patients with higher HBcrAg levels. Further studies with a larger cohort are needed, as our study included relatively few patients with very high HBsAg and HBcrAg levels.

Our study included a diverse cohort of both treated and treatment-naïve patients. The reduced response to DAC/αPD-L1 observed in treated patients may not be directly attributable to the therapy itself but could be influenced by other factors, such as the initial disease stage prior to treatment. In the treated patients, therapy was initiated due to active disease distinguishing them from the predominantly inactive carriers among the treatment- naïve patients [34]. Given the diverse stages of chronic HBV infection and therapy statuses in our study, the patients might have distinct epigenetic signatures that are differentially affected by DAC treatment. Usually, DNA methylation signatures are considered stable in the short-term and can even be passed on to subsequent generations [35]. However, environmental factors and life style can still influence the imprints [36]. Whether NA- treatment and subsequent viral suppression influence the epigenetic immune signature remains elusive.

We observed HBV epitope-specific CD8^+^ T cell response differences to DAC, aligning with recent findings on variability in HBV core and pol-specific CD8^+^ T cell responses [24,37–39]. Specifically, pol_455_-specific CD8^+^ T cells showed enhanced responses to the DAC/αPD-L1 combination compared to αPD-L1 alone, whereas core_18_-stimulated T cells exhibited more heterogeneous responses. Previous studies have demonstrated that pol_455_-specific CD8^+^ T cells are more impaired than core_18_-specific T cells [24,37–39], which may explain the more pronounced effect of DAC/αPD-L1 on pol_455_-specific CD8^+^ T cells.

This study supports previous research on chronic infections [5,18,40,41] by highlighting the persistent epigenetic imprint of exhaustion that can impede HBV-specific immune responses and underscores the potential of epigenetic-modifying approaches. The DAC/αPD-L1 combination notably enhances the immune response, particularly in more exhausted pol455-specific CD8^+^ T cells. Given their documented phenotypic and metabolic differences [24,37–39], epitope-specific variations in epigenetic signatures may contribute to the observed differences in response to DAC/αPD-L1 treatment.

Due to the very small number of HBV-specific cells and the high cell requirement of current epigenetic methods, methylation analysis was conducted on bulk PBMCs and not on antigen-specific T cells. This represents a limitation of the study, as this may obscure DNA methylation signatures specific to exhausted T cells, reducing the applicability of the bulk epigenetic profile to these cells and complicating the analysis of epitope-specific imprints. Despite this limitation, we observed significant differences in methylation levels between DAC-responders and non-responders, particularly in the *IFNG* gene, which was also reflected by *ex vivo* IFNγ plasma levels.

A further limitation of this study is the non-specificity of both checkpoint inhibition and DAC treatment. Unlike targeted therapies, the broad effect of DAC on DNA methylation patterns may lead to variations in treatment response among patients and it remains unclear which additional immune cells might be demethylated. Moreover, DAC may not only inhibit the methylation of exhaustion-associated genes but also influence other hypermethylated genes affecting treatment response. According to Klco et al. [32], DAC-induced hypomethylation mainly occurs in hypermethylated regions which might explain that the most substantial methylation differences between DAC-responders and non-responders were observed in genes that were initially more hypermethylated. Further investigation is needed to determine whether hypermethylation is a general effect of chronic infection or inflammation or specific to HBV. Nonetheless, the enhanced response in more exhausted pol_455_-specific T cells highlights a virus-specific aspect of the treatment’s efficacy. In line with this, a recent tumor mouse study suggested that DAC monotreatment might demethylate nonspecifically, but the combination with checkpoint inhibitor precisely promoted the expansion of tumor-specific progenitor exhausted T cells [42]. Furthermore, genetic factors may also play a role in responses to DAC, as documented in an AML study [43], where mutations in the *DNMT3A* gene influenced sensitivity to hypomethylating agents like DAC, which have not been investigated in this study.

In conclusion, our study highlights the important role of immune epigenetics in chronic HBV infection and its impact on the efficacy of immunomodulatory treatments such as αPD-L1. We propose that epigenetic modulation, particularly with the DNA methyltransferase inhibitor, DAC, is a promising strategy to improve the immune response in chronic HBV infection, a potential supported by previous data from the LCMV *in vivo* mouse model. However, the observed heterogeneity of patient responses underscores the complexity of host-virus interactions and the need for personalized treatment approaches. While further investigation of the underlying mechanisms is needed, our results provide valuable insights into the potential of a combination of epigenetic modulation and immune checkpoint blockade as a pathway to a functional cure of HBV infection.

## Acknowledgements

Funded by the German Research Foundation (Deutsche Forschungsgemeinschaft, DFG) under Germany’s Excellence Strategy - EXC 2155 - project number 390874280, and by the German Center for Infection Research (Deutsches Zentrum für Infektionsforschung, DZIF) – TTU 05.820 and TTU 05.708. MUQ, KS, and YC were supported by the Hannover Biomedical Research School (HBRS) and the Center for Infection Biology (ZIB). MUQ presented the data at the EASL Congress 2023, GASL 2023 and DZIF TTU Hepatitis Meeting 2024.

## Author contributions

MUQ, AK, and MC designed the study. MUQ, KS, and YC conducted experiments. MUQ, BB, HS, KS, and YC acquired data. MUQ, KS, and YC analyzed the *in vitro* data. MUQ, and MG analyzed the epigenetic data. MUQ, KS, MG, CX, AK, and MC interpreted the data. HW, BM, and KD provided essential patient samples and clinical data. MUQ, YC, AK and MC prepared the manuscript. All authors provided critical review of the manuscript.

## Abbreviations

ALT: alanine transaminase
αPD-L1: anti-programmed cell death ligand-1 antibody
AST: aspartate transferase
CHB: chronic HBV
CI: confidence interval
DAC: decitabine
HBcrAg: hepatitis B core- related antigen
HBeAg: hepatitis B e antigen
HBsAg: hepatitis B surface antigen
HBV: hepatitis B virus
IFNγ (*IFNG*): interferon gamma
NA: nucleos(t)ide analogues
neg: negative
OLP: overlapping peptide pool
PBMC: peripheral blood mononuclear cell
PD-1 (*PDCD1*): programmed cell death 1
pol: polymerase
pos: positive
TNFα: tumour necrosis factor alpha
TSS: transcription start site.

