## Supplemental methods/tables/figures for "Enhancing HBV-specific T cell responses through a combination of epigenetic modulation and immune checkpoint inhibition"

***Modulation of HBV-specific T cell response after in vitro stimulation with HBV-specific peptides***

HBV-specific T cell responses were analyzed as previously described [1]. Briefly, thawed PBMCs were stimulated for 10 days with either HBV core OLP or HLA-A*02-restricted HBV-specific peptides, $core_{18}$ and $pol_{455}$.

For the 10-day stimulation, $3\times10^5$ cells/well were resuspended in AB-medium (10 % human AB serum, 1 % non-essential amino acids, 1 % sodium pyruvate, 1 % penicillin/streptomycin, 5 mM HEPES buffer, 1 mM L-glutamine in RPMI-1640 medium with GutaMAX, (Gibco, CA, USA)) and seeded into 96-well round bottom plates. The PBMCs were stimulated either with HBV OLP pool or HLA-A*02-restricted peptides at day 0. Besides peptide-treated wells, a negative medium control (unstimulated and untreated background control) and a well for phorbol-12-myristate-13-acetate (PMA)/ionomycin as a positive control were included. As treatments, different combinations of DAC with or without checkpoint inhibitor αPD-L1 (10 µM, clone M1H1, Invitrogen, CA, USA) were used. For initial experiments, αPD-L1 was added simultaneously with DAC at day 0. Later, we decided to continue with the DAC pretreatment approach and αPD-L1 addition at day 3. Therefore, three days after stimulation, the cells were treated with or without the addition of αPD-L1. At day 3 and 7 post-stimulation, cells were fed with recombinant human IL2-containing medium (5 IU/mL, Peprotech, NJ, USA). After 10 days, the cells were restimulated with the respective HBV OLP pools/HLA-A*02-restricted peptide with or without αPD-L1. After 1 hour (h) incubation at 37 °C, the cells were treated with Brefeldin A (2 µg/mL, Sigma-Aldrich, Germany). After another 5 h incubation, the cells were washed with FACS buffer and stained extracellular with antibodies (Supplementary Table 1), dead cell marker (DCM, LIVE/DEAD Aqua cell stain kit, Life Technologies, USA, CA) and Fc-receptor blocking reagent for 15 min in the dark at 4 °C. After the staining, the cells were washed with DPBS and afterwards resuspended with Foxp3/Transcription Factor Staining Buffer Set (eBioscience, Germany, Frankfurt) for 30 minutes fixation at 4 °C. The fixation was stopped by washing the cells twice with permeabilization buffer before the cells were stored in the buffer in the dark overnight at 4 °C. The next day, the cells were stained intracellular (Supplementary Table 1) for 30 min at 4 °C, then washed twice in permeabilization buffer and subsequently resuspended in FACS buffer for data acquisition on a BD LSRFortessa flow cytometer (BD Biosciences, Germany, Heidelberg). Flow cytometry data were analyzed using FlowJo software (V.10.8.1).

### Supplementary tables

#### *Supplementary Table 1: Overview of used antibodies and viability stains.*

| Marker | Clone | Fluorochrome | Manufacturer | Catalog number |
|---|---|---|---|---|
| CD14 | M5E2 | V500 | BD Biosciences | 561391 |
| CD19 | SJ25C1 | BV510 | BD Biosciences | 562947 |
| CD3 | UCHT1 | PE-Cy5 | Biolegend | 300410 |
| CD4 | RPA-T4 | BB515 | BD Biosciences | 564419 |
| CD8 | RPA-T8 | BV605 | Biolegend | 301040 |
| Granzyme B | GB11 | Alexa Fluor 647 | Biolegend | 561999 |
| HLA-DR | L243 | BV785 | Biolegend | 307642 |
| IFNγ | B27 | BV421 | BD Biosciences | 562988 |
| LIVE/DEAD™ Fixable Aqua Dead Cell Stain Kit | - | Aquamarine | Thermo Fisher Scientific | L34957 |
| PD-1 | EH12.1 | PE | BD Biosciences | 560795 |
| TNFα | MAb11 | BV650 | Biolegend | 502938 |
| Mip-1β | D21-1351 | PE-Cy7 | BD Biosciences | 560687 |
| Ki67 | Ki-67 | BV711 | Biolegend | 350515 |

### Supplementary figures

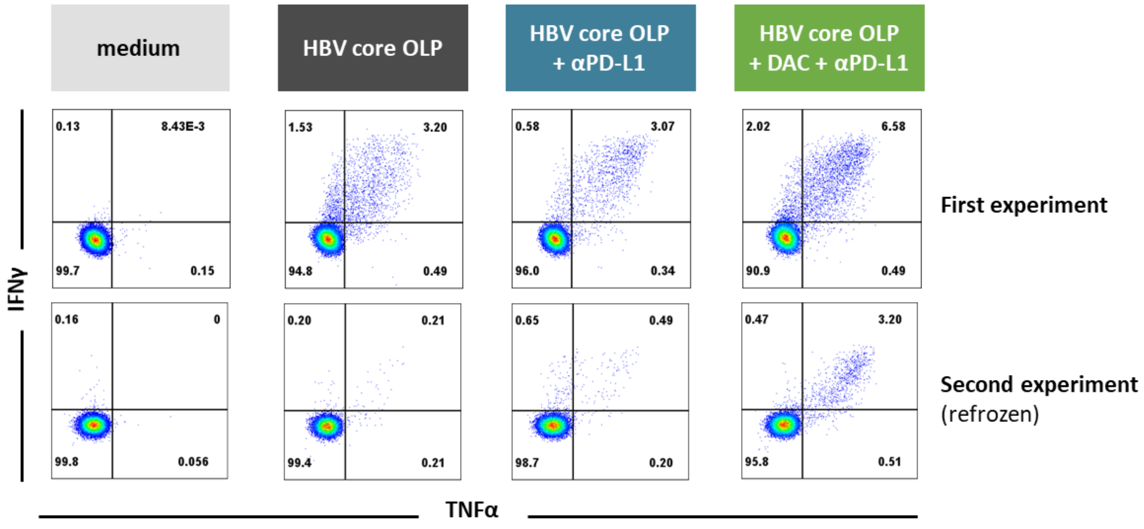

**Supplementary Figure 1: Repetition of 10-day culture of a refrozen sample.** Flow cytometry plots of one sample that was refrozen and used in a second independently performed 10-day culture experiment. Abbreviations: αPD-L1, anti-programmed cell death ligand-1 antibody; DAC, decitabine; HBV, hepatitis B virus; IFNγ, interferon gamma; OLP, overlapping peptide pool; TNFα, tumour necrosis factor alpha.

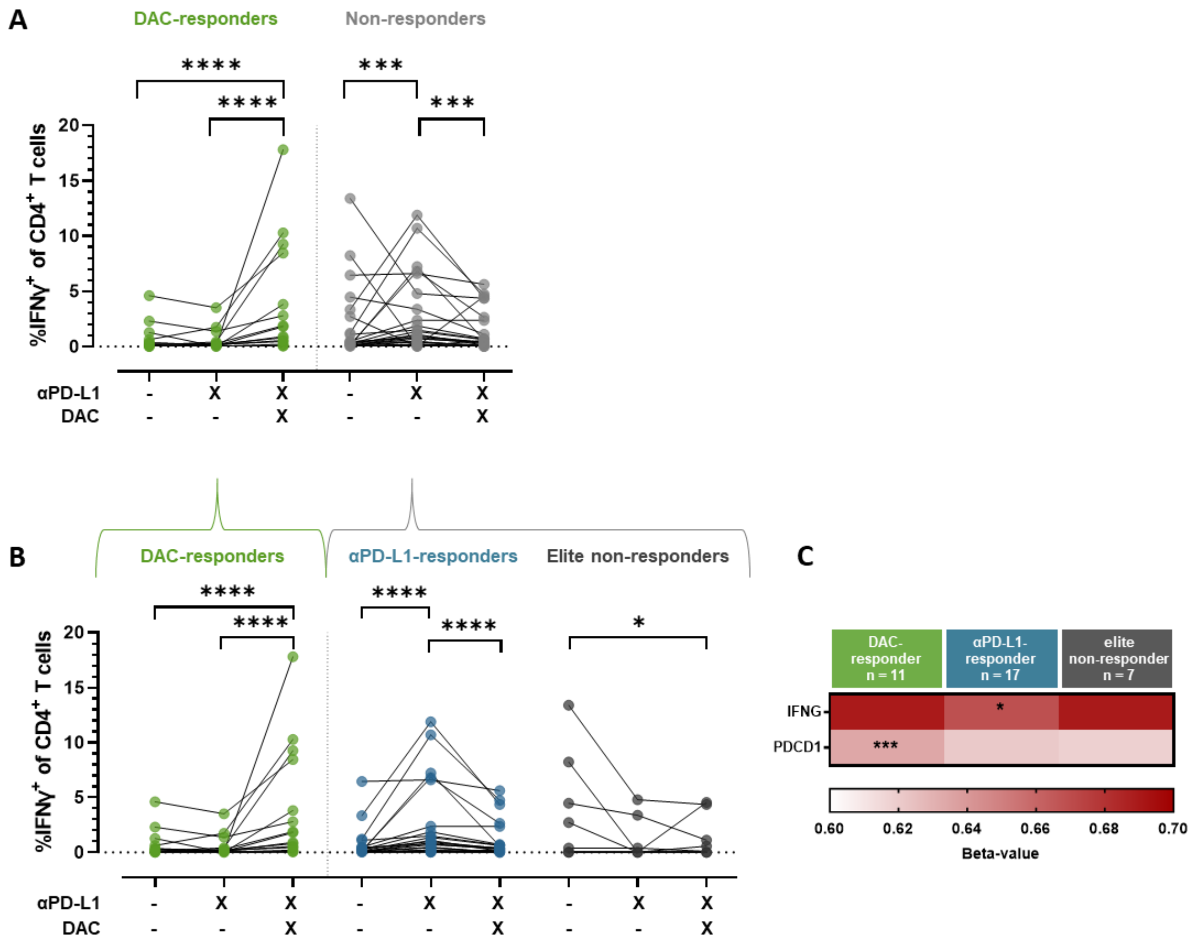

**Supplementary Figure 2: Comparative analysis of immune responses of DAC-responders and non-responders.** Frequency of IFNγ$^+$ CD4$^+$ T cells following a 10-day culture as depicted in Figure 2D separated either into (**A**) DAC-responders (green, n = 17) and non-responders (grey, n = 36) or (**B**) DAC-responders (n = 17) versus αPD-L1-responders (blue, n = 24) and elite non-responders (dark grey, n = 12). (C) Mean methylation of gene-associated CpG sites of DAC-responders (n = 11), αPD-L1-responders (n = 17), and elite non-responders (n = 7). Statistical significances were determined using the Friedman test with Dunn's multiple comparison (p*<0.05; p***<0.001; ****p<0.0001). Statistically significant different group in C was marked. Abbreviations: αPD-L1, anti-programmed cell death ligand-1 antibody; DAC, decitabine; IFNγ, interferon gamma.
